# Methotrexate carried in lipid core nanoparticles reduces microglial activation and is neuroprotective after ischemic cortical stroke

**DOI:** 10.1101/2020.06.16.155804

**Authors:** Edmundo L. R. Pereira, Michelle N.C. Dias, Ijair R. dos Santos, Ana Carolina Ramos, Moisés Hamoy, Danielle Cristine A. Feio, Thauany M. Tavoni, Priscila Carvalho M. da Silva, Raul Maranhão, Walace Gomes-Leal

**Author notes:** Corresponding Author: Dr. Walace Gomes-Leal, Laboratory of Experimental Neuroprotection and Neuroregeneration, Institute of Biological Sciences, Federal University of Pará. Rua Augusto Corrêa S/N. Campus do Guamá, CEP: 66075-900. Belém-Pará Brasil.

## Abstract

Methotrexate carried in lipid core nanoparticles (LDE-MTX) is a low toxicity compound effective in reducing inflammation and secondary damage in experimental models of arthritis, atherosclerosis, myocardial infarction, cardiac allograft vasculopathy and other pathological conditions. Nevertheless, whether it is neuroprotective after stroke is unknown. Here, we explored whether LDE-MTX could cross blood brain barrier (BBB) to exert anti-inflammatory and neuroprotecive effects after experimental cortical stroke in rats. Tissue uptake was assessed by injecting radioactively labeled-LDE through the caudal vein into both sham (n=18) and adult Wistar rats submitted to endothelin-1 (ET-1)-induced cortical stroke (n=11). To address possible neuroprotective effects of LDE-MTX after stroke, 10 adult male Wistar rats were randomly allocated in two groups: animals treated with LDE-MTX (1 mg/kg, *i.v.*, n=5) or LDE-alone (*i.v.*, n=5) at 4 hours after stroke induction. Animals were perfused with 0.9% saline and 4% paraformaldehyde at 7 days post-injury. Histopathology was assessed by cresyl violet staining. Mature neuronal bodies (anti-NeuN), astrocytes (anti-GFAP) and microglia (anti-Iba1) were immunolabeled by immunohistochemistry. Scintigraphy technique revealed accumulation of tritiated LDE in different brain regions and in non-neural organs without overt toxicity in both sham and ischemic rats. LDE-MTX treatment induced a 10-fold (1000%) reduction in microglial activation in the ischemic cortex and afforded a 319% increase in neuronal preservation in the ischemic periinfarct region compared to LDE-alone group. There was no effect of LDE-MTX treatment on primary infarct area and astrocytosis. The results suggest that LDE-MTX formulation must be considered a very promising neuroprotective agent for ischemic stroke. Future studies using different concentrations and longer survival times are needed before assessing the suitability of LDE-MTX as a neuroprotective agent for human stroke.

## 1. Introduction

Brain stroke is a major public health problem is several countries and a devastating acute neural disorder caused by obstruction (ischemic) or rupture (hemorrhagic) of blood vessels (kaiser & West, 2020). Following metabolic collapse with subsequent paucity of glucose and oxygen, a primary damage occurs followed by a secondary damage caused by a myriad of pathological events, including spread depression, excitotoxicity, neuroinflammation, oxidative stress, programmed cell death, metabolic acidosis (Lo, 2003; Moskowitz *et al.*, 2010).

In the case of ischemic stroke, the lesion center may expand over the ischemic penumbra causing a net expansion of more than 65 % in comparison to the initial primary damage (Lo, 2003; Moskowitz *et al.*, 2010). The main purpose of experimental neuropathology is to develop neuroprotective agents to avoid secondary damage (Cook *et al.*, 2011; kaiser & West, 2020) or contributes to repair by cell replacement and other neuroplasticity mechanism achieving functional recovery (Boltze et al., 2019; Palma-Tortosa et al., 2020). This has been challenging considering the diversity of the damaging mechanisms, inadequacy of experimental models to simulate human stroke and great rate of fail of tested drugs (Dirnagl, 2006).

It has been established that neuroinflammation plays a pivotal role on the pathophysiology of acute (Gomes-Leal, 2012; Gomes-Leal, 2019) and chronic (Ransohoff, 2016) neural disorders. After stroke, an intense microglial activation takes place lasting several weeks after primary damage (Thored *et al.*, 2009) with both beneficial and detrimental consequences (Gomes-Leal, 2012; Gomes-Leal, 2019).

Inhibition of microglial activation with the semi-synthetic tetracycline minocycline reduces neuroinflammation and infarct area in both cortex and striatum after middle cerebral artery occlusion (MCAO) in adult rats (Yrjanheikki *et al.*, 1999). Considerable neuroprotection has also been reported in the hippocampal CA1 region following inhibition of microglial activation using PJ34, a Poly(ADP-ribose) polymerase inhibitor after forebrain ischemia (Hamby *et al.*, 2007). We have previously shown that modulation of microglia activation with minocycline improves therapeutic effects of bone marrow mononuclear cells after both cortical (Franco *et al.*, 2012) and striatal (Cardoso *et al.*, 2013) stroke.

Microglial response may also be beneficial after stroke and other central nervous system (CNS) disorders (Gomes-Leal, 2012; Gomes-Leal, 2019). Genetic ablation of proliferating microglia exacerbates neuroinflammation, increases programmed cell death and enlarges infarct area following MCAO (Lalancette-Hebert *et al.*, 2007). Inhibition of microglial activation with minocycline increases neuronal death following oxygen-glucose deprivation in organotypic hippocampal slice culture (Neumann *et al.*, 2006a) and microglia are beneficial by engulfment of neutrophils in this same experimental of model of ischemia (Neumann *et al.*, 2008). We have recently hypothesized that microglia may be beneficial or contribute to neuronal death depending on the nature of danger signals present over the pathological environment – the friendly fire hypothesis (Gomes-Leal, 2012; Gomes-Leal, 2019).

The search for new neuroprotective agents for stroke may rely on the nanotechnology development. Nanomedicine applications include the use of nanoparticles with specific physical and chemical properties, which serve as vehicles for a particular therapeutic agent. Liposomes are especially suitable for these purpose, because they possess structure that resembles cholesterol-carrying particles such as low density lipoprotein (LDL), which assures that the therapeutic agent reaches in a more efficacious way the pathological environment (Bobo *et al.*, 2016).

We have developed an emulsion lipid system composed of nanoparticles of medium size 30 nm (LDE) functionally similar to low density lipoprotein (LDL), but with different protein composition, which allows more efficient internalization into the cells (Maranhao *et al.*, 1993). LDE has been used as an efficient carrier for the transport of anti-inflammatory and chemotherapeutic substances, including docetaxel (Meneghini *et al.*, 2019), carmustine (Daminelli *et al.*, 2016), paclitaxel (Maranhão et al., 2008) and methotrexate (MTX) (Bulgarelli et al., 2013; Mello et al., 2013; Maranhao *et al.*, 2017) in different experimental models of non-neural diseases and even in a clinical trial for epithelial ovarian carcinoma in humans (Graziani *et al.*, 2017).

Commercial MTX is widely used as an anti-inflammatory agent in rheumatic diseases and we found that association with LDE increased ninetyfold the uptake of the drug by cells (Moura et al., 2011). Subsequently, it was shown that LDE-MTX had the ability to markedly reduce the area of atherosclerotic lesions of cholesterol-fed rabbits and to decrease the intimal area, specifically the invasion of macrophage and smooth cells (Bulgarelli et al., 2013).

More recently, rats with acute myocardial infarction induced by ligation of the left coronary artery treated with LDE-MTX display a remarkable improvement of the heart function and reduction of the infarction size (Maranhão et al., 2017). In contrast, the effects of commercial MTX were minor and did not improve the heart function or the infarct size. Taken together, the results from those studies prompted us to test the hypothesis that LDE-MTX could be neuroprotective following experimental stroke.

In the present study, we first tested the possibility that intravenously infused LDE-MTX may cross the blood brain barrier (BBB) in both sham and ischemic rats. Second, we explored whether LDE-MTX available in the brain parenchyma may be neuroprotective after cortical ischemia in adult rats.

## 2. Material and methods

### 2.1. Animals

Male adult Wistar rats (250-300 g) were obtained from the Federal University of Pará Central Animal Facility. All animals were housed under standard conditions with food and water available ad libitum. All experimental procedures were carried out in accordance with the Principles of Laboratory Animal Care (NIH publication No. 86-23, revised 1985) and European Commission Directive 86/609/EEC for animal experiments under license of the Ethics Committee on Experimental Animals of the Federal University of Pará. All possible efforts were made to avoid animal suffering and distress.

### 2.2. Experimental model of cortical ischemia

Focal cortical ischemia was induced by microinjections of endothelin-1 (ET-1, (Sigma, Saint Louis, MO, USA)-a powerful vasoconstrictor whose brain microinjections cause considerable reduction of blood flow (Fuxe *et al.*, 1989; Agnati *et al.*, 1991). We have previously validated the ET-1 model of stroke in several investigations (Dos Santos *et al.*, 2007; Franco *et al.*, 2012; Cardoso *et al.*, 2013; Lopes *et al.*, 2016; Lima et al., 2016; Souza *et al.*, 2017).

In short, animals were anesthetized with ketamine hydrochloride (72 mg/kg, *i.p*) and xylazine hydrochloride (9 mg/kg, *i.p*) and held in a stereotaxic frame (Insight, Brazil) after their corneal and paw withdraw reflexes were abolished. A homoeothermic blanket unit was used to maintain animal’s body temperature, as measured by a rectal thermometer. After craniotomy, 40 pmol of ET-1 (Sigma, Saint Louis, MO, USA) in l μ1 of sterile saline was injected into the rat motor cortex (n=5 per survival time/animal group) over a period of 2 minutes using a finely drawn glass capillary needle. The capillary needle was left in position for 3 minutes before being slowly withdrawn. Control animals were injected with the same volume of sterile saline (n=5 per survival time). We used the following stereotaxic coordinates in relation to the bregma: +2.3 mm lateral; +1.2 mm posterior and 0.50 mm deep from the pial surface in the dorsoventral axis (Paxinos *et al.*, 1980). To identify the injection site, a small quantity of colanyl blue was added to both ET-1 and vehicle solutions. After surgery, animals were allowed to recover with free access to food and water for 7 days.

### 2.3. Animals groups and treatment

Experiments were performed in two steps. First, to assess whether LDE crosses the BBB, we delineated the following experimental groups

a1. Sham group (n=18): animals without ischemia treated with 200 µL (480,000 cpm) of LDE labeled with 3H-cholesteryl oleate ether (3H-LDE) intravenously injected through the caudal vein. Animals were anesthetized with ketamine hydrochloride (72 mg/kg, *i.p*) and xylazine hydrochloride (9 mg/kg, *i.p*) and perfused with 0.9% sterile saline without perfusion at 7 days post-injection;

b1. Ischemic group (n=11): ischemic animals treated with 200 µL (480,000 cpm) of 3H^+^-LDE intravenously injected through the caudal vein. Brains were removed as described for sham animals.

In the second phase of this study, we explored the potential of LDE-MTX as a neuroprotective agent. For this purpose, the following experimental groups were delineated:

a2. LDE-MTX (n=5): ischemic animals treated with LDE-MTX (1 mg/Kg) intravenously injected through caudal vein at 4 hours after stroke induction. Animals were perfused with 0.9% sterile saline and 4% paraformaldehyde at 7 days post-injury.

b2. Control group (n=5): ischemic animals treated with unconjugated LDE (1 mg/Kg) intravenously injected through caudal vein at 4 hours after stroke induction. Animals were perfused with 0.9% sterile saline and 4% paraformaldehyde at 7 days post-injury.

### 2.4. LDE preparation and association with MTX

LDE is a lipid core nanoparticle that was developed by Professor Raul C. Maranhão at the Laboratory of Metabolism and Lipids, Heart Institute, Faculty of Medicine, University of São Paulo, Brazil (InCor/FMUSP) (Maranhãoet al., 1993).

LDE preparation consists of a lipid mixture composed of 100 mg cholesteryl oleate, 200 mg egg phosphatidylcholine (Lipoid, Germany), 10 mg triglycerides, 12 mg cholesterol and 60 mg of derivatized MTX according to a previously described protocol (Maranhão et al., 2017). The aqueous phase (100 mg of polysorbate 80 and 10 mL of Tris-HCl buffer, pH 8.05) was kept at room temperature and the pre-emulsion obtained by adding the hydrophilic phase to the oil phase by ultrasonic radiation, until complete dissolution of the drug.

Emulsification of the compounds was obtained by high-pressure homogenization using an Emulsiflex C5 homogenizer (Avestin, Canada). After homogenization at constant temperature, the nanoemulsion was centrifuged at 1800 g for 15 min at 4°C to separate the emulsified product from unbound MTX. The particle size of preparations was 60 nm, as measured by dynamic light scattering method at a 90° angle, using the ZetaSizer Nano ZS90 equipment (Malvern Instruments, Malvern, UK). The efficiency of the association of MTX to LDE was measured by HPLC. The nanoparticles were sterilized by passing through a 0.22 μm pore polycarbonate filter (EMD Millipore Corporation, Billerica, MA, USA) and kept at 4 °C until it was used.

### 2.5. LDE uptake by the brain parenchyma

For LDE tissue uptake, the same LDE preparation was used, except that the lipids were labelled with [3H]-cholesteryl oleate ether (Perkin Elmer, Boston, MA, USA) and without MTX addition. 3H-LDE was administered through the rat caudal vein in a single standardized dose (200 µL, 480,000 cpm). Animals from Sham and ischemic groups were perfused with 0.9% sterile saline only, without paraformaldehyde, and brain (neocortex from both hemispheres, cerebellum, brain stem), muscle (right rectus abdominis) and liver samples were collected.

Tissues samples were kept in cold saline solution before lipid extraction with chloroform/methanol (2:1 v/v) (Folch et al., 1957). After extraction, the solvent was evaporated under N2 flow and resuspended with 500 mL of chloroform/methanol (2:1v/ v) and half the suspension was placed separately into vials with 5 mL of scintillation solution (Ultima Gold XR, Perkin Elmer). Levels of radioactivity were measured on the Liquid Scintillation Analyzer, 1600TR Tri-Carb, Packard (Palo Alto, CA, USA).

### 2.6. LDE-MTXc treatment

In order to investigate whether LDE-MTX has neuroprotective and anti-inflammatory effects after cortical ischemia, animals from second phase of the study were submitted to ET-1 microinjections according to a stroke model routinely used in our laboratory (Dos Santos *et al.*, 2007; Franco *et al.*, 2012; Cardoso *et al.*, 2013; Lopes *et al.*, 2016; Souza *et al.*, 2017). Four hours after ischemic induction, animals were intravenously treated LDE-MTX or unconjugated LDE in the concentration of 1 mg/kg. This LDE-MTX dose afforded considerable anti-inflammatory and tissue protective effects in experimental models of arthritis (Mello *et al.*, 2013) and myocardial infarction (Maranhao *et al.*, 2017) in our previous studies.

Animals from both experimental groups were perfused at 7 days post stroke onset, as tissue damage and neuroinflammation are maximum at this survival time (Morioka *et al.*, 1993; Lima *et al.*, 2016).

### 2.7. Tissue processing and perfusion

After 7 days post-injury, animals were deeply anesthetized (*i.p*) with a mixture of ketamine hydrochloride (72 mg/Kg) and xylazine (9 mg/kg). After absence of both corneal and paw withdraw reflexes, animals were perfused with 0.9% saline solution followed by 4% paraformaldehyde. Brains were removed and post-fixed for 24 hours in the same fixative used in the perfusion and cryoprotected in gradients of a solution containing glycerol and sucrosis. They were then embedded in Tissue Tek, frozen at – 55 °C degrees in a cryostat chamber (Carl Zeiss, Micron, Germany), and cut at 30 μm thickness. 50 μm coronal sections were also obtained for gross histopathological analysis using cresyl violet staining (Lima *et al.*, 2016). Sections were directly collected onto gelatinized slides for better adherence and kept at room temperature for at least 24 hours until being kept in a freezer at −20 °C.

### 2.8. Gross histopathology

The lesion area was assessed in 50 μlm sections stained by cresy violet (Sigma, Saint Louis, MO, USA). Infarct size was recognized by the presence of colanyl blue, pallor, inflammatory infiltrate and necrosis as previously described (Franco *et al.*, 2012).

### 2.9. Immunohistochemistry

We have previously determined several neuropathological parameters to assess neuroprotection in different experimental models of acute brain (Guimaraes *et al.*, 2010; Franco *et al.*, 2012; Guimaraes-Santos *et al.*, 2012; Cardoso *et al.*, 2013; Lima *et al.*, 2016; Lopes *et al.*, 2016) and spinal cord injuries (Gomes-Leal *et al.*, 2004; Gomes-Leal *et al.*, 2005).

To investigate whether LDE-MTX modulates neuroinflammation and is neuroprotective after cortical ischemia, we used the following antibodies: anti-NeuN (1: 100, Chemicon) as a marker for mature neurons. It recognizes a specific epitope in the nucleus of differentiated neurons (Mullen *et al.*, 1992); Anti-Iba1 (1: 1000, Wako) as a specific microglial marker. It labels a calcium-binding protein present in the cytoplasm of microglia (Ito *et al.*, 1998); anti-ED-1 (Serotec, 1: 200). It labels an epitope on the lysosome membrane of activated macrophages/microglia (Dijkstra *et al.*, 1985); anti-glial acid fibrillary protein (GFAP) (Dako, 1: 1000). A classical astrocyte marker (Gomes-Leal *et al.*, 2004).

Details on the immunolabeling were described in our previous papers (Gomes-Leal *et al.*, 2004; Franco *et al.*, 2012). In short, mounted sections onto gelatinized slides were removed from the freezer, dried in an oven for 30 minutes at 37°C, washed in PBS (phosphate buffer saline) under constant agitation for 5 minutes and immersed into borate buffer (0.2M, pH 9.0) at 65 °C for 20 minutes. Sections were then cooled in the same solution for the same period of time at room temperature. Sections were then washed three times in PBS for 5 minutes and immersed in methanol containing hydrogen peroxide (1ml/100ml of methanol).

After antigen retrieval, sections were washed 3 times in PBS/Tween (5 minutes each) and incubated in 10% normal goat serum (anti-Iba1 and anti-GFAP) or normal horse serum (anti-ED1 and anti-NeuN) for 1 hour. After this period, they were incubated in primary antibody diluted in 10% normal serum overnight using the previously mentioned dilution. In the next day, sections were washed again (3 times in PBS/Tween, 5 minutes each) and incubated in biotinylated goat anti-rabbit (1: 200, Vector Laboratories, USA) or biotinylated horse anti-mouse (1: 100, Vector Laboratories, USA) diluted in PBS for 2 hours.

One hour before incubation in the secondary antibody, ABC solution (avidin-biotin-peroxidase-ABC Elite kit, Vector laboratories, USA) was prepared and allowed to rest at least for 30 minutes with no agitation. After further washing, sections were incubated in ABC solution diluted in PBS for 2 hours. They were then washed four times and stained using diaminobenzidine (DAB, Sigma-Aldrich, USA). Subsequently, sections were washed three times in 0.1M PB, dehydrated in alcohol gradients, cleared in xylene and coversliped using Entellan (Merk, USA).

### 2.10. Qualitative analysis

All sections stained with the different histological methods were observed using a light microscopy (Nikon Eclipse 50i, Nikon, Tokyo, Japan). High resolution images from the more representative fields were obtained using a digital camera (Moticam 2500) attached to a light microscope (Nikon Eclipse 50i, Nikon,Tokyo, Japan).

### 2.11. Quantitative analysis

Numbers of NeuN+ cells and activated macrophages/microglia (Iba1+ cells) cells were counted using a square 0.25 mm-wide grid (objective 40x) in the eyepiece of a microscope (Nikon-Eclipse 50i, Japan) for all experimental groups. This grid corresponds to an area of 0.0625 mm^2^. 3 sections per animal, 3-4 fields per section, with five animals per survival time, were used in the counts. For NeuN + cells, four counting fields in both lateral and medial part of ischemic core, were used.

In the quantitative analysis of Iba1+ cells, three non-overlapping fields were counted in the ischemic core, in which the largest numbers of phagocytes are present, for all experimental animals. Only rounded (phagocytes) Iba1 + cells were counted according to protocol previously reported by our group (Franco *et al.*, 2012). This procedure was performed to quantitatively assess the effect of LDE-MTX on the maximum morphological level of microglial activation (Thored *et al.*, 2009; Franco *et al.*, 2012).

### 2.12. Statistical analysis

Normality test was applied to the data. Descriptive statistic was performed for all counts. Averages, standard deviations, and standard errors were calculated. Comparisons between groups were assessed by unpaired Student’s *t* test. Statistical significance level was accepted at *p*<0.01. All statistical analyses were performed using the Software GraphPad Prism 7.0.

For data obtained from LDE tissue uptake, the following non-parametric tests were applied: Mann-Whitney and Kruskal-Willis tests. Statistical significance level was accepted at p<0.01. All statistical analyses were performed using the Software GraphPad Prism 7.0.

## 3. Results

### 3.1 3H-LDE *crosses blood brain barrier and accumulates in different brain regions*

In the first phase of the study, we used the scintigraphy technique to assess whether intravenously injected 3H-LDE crosses the BBB in sham or ischemic adult rats. 3H-LDE nanoparticles were found in samples of the brain, muscle and liver of all investigated animals (Table 1). There was, as expected, preponderance of particle uptake by the liver, while the lowest values were present in the brain and rectus abdominis muscle (Tables 1 and 2, Fig. 1).

**Table 1.** Scintigraphy data for tissue uptake of [3H]-cholesteryl oleate ether-LDE in Sham and ischemic animals (cpmb/g tissue)

| *Animal* | *ET-1* | *Brain* | *Muscle* | *Liver* |
| --- | --- | --- | --- | --- |
| 1 | + | 13 | 7 | 550 |
| 2 | + | 31 | 11 | 812 |
| 3 | + | 7 | 27 | 652 |
| 4 | + | 17 | 24 | 719 |
| 5 | + | 55 | 22 | 2423 |
| 6 | + | 18 | 25 | 1578 |
| 7 | + | 20 | 31 | 2159 |
| 8 | + | 27 | 18 | 668 |
| 9 | + | 19 | 21 | 827 |
| 10 | + | 74 | 26 | 442 |
| 11 | + | 20 | 68 | 2540 |
| 12 | - | 35 | 144 | 9450 |
| 13 | - | 36 | 37 | 9926 |
| 14 | - | 74 | 137 | 9789 |
| 15 | - | 35 | 116 | 9847 |
| 16 | - | 61 | 132 | 9807 |
| 17 | - | 15 | 154 | 8304 |
| 18 | - | 80 | 55 | 9864 |
| 19 | - | 77 | 156 | 9767 |
| 20 | - | 33 | 29 | 9957 |
| 21 | - | 14 | 517 | 9469 |
| 22 | - | 41 | 81 | 1779 |
| 23 | - | 16 | 250 | 934 |
| 25 | - | 19 | 42 | 3035 |
| 26 | - | 10 | 283 | 1004 |
| 25 | - | 12 | 154 | 9574 |
| 27 | - | 18 | 162 | 9247 |
| 28 | - | 38 | 354 | 8681 |
| 29 | - | 42 | 20 | 2238 |

ET-1 = endothelin-1

**Table 2.** Normalized percentage of tissue uptake of [3H]-cholesteryl oleate ether-LDE in sham and ischemic animals (%/g tissue).

| *Tissue* | *Sham* | *Stroke* |
| --- | --- | --- |
| *Brain* | 0.671 ± 0.134 | 2.012 ± 0.342* |
| *Muscle* | 3.969 ± 1.521 | 2.097 ± 0.311 |
| *Liver* | 95.360 ± 1.561 | 95.891 ± 0.415 |

Data are expressed as mean ± SEM. \* $p=0.0038$ (Unpaired *Student t* test with Welch correction test).

**Figure 1.**
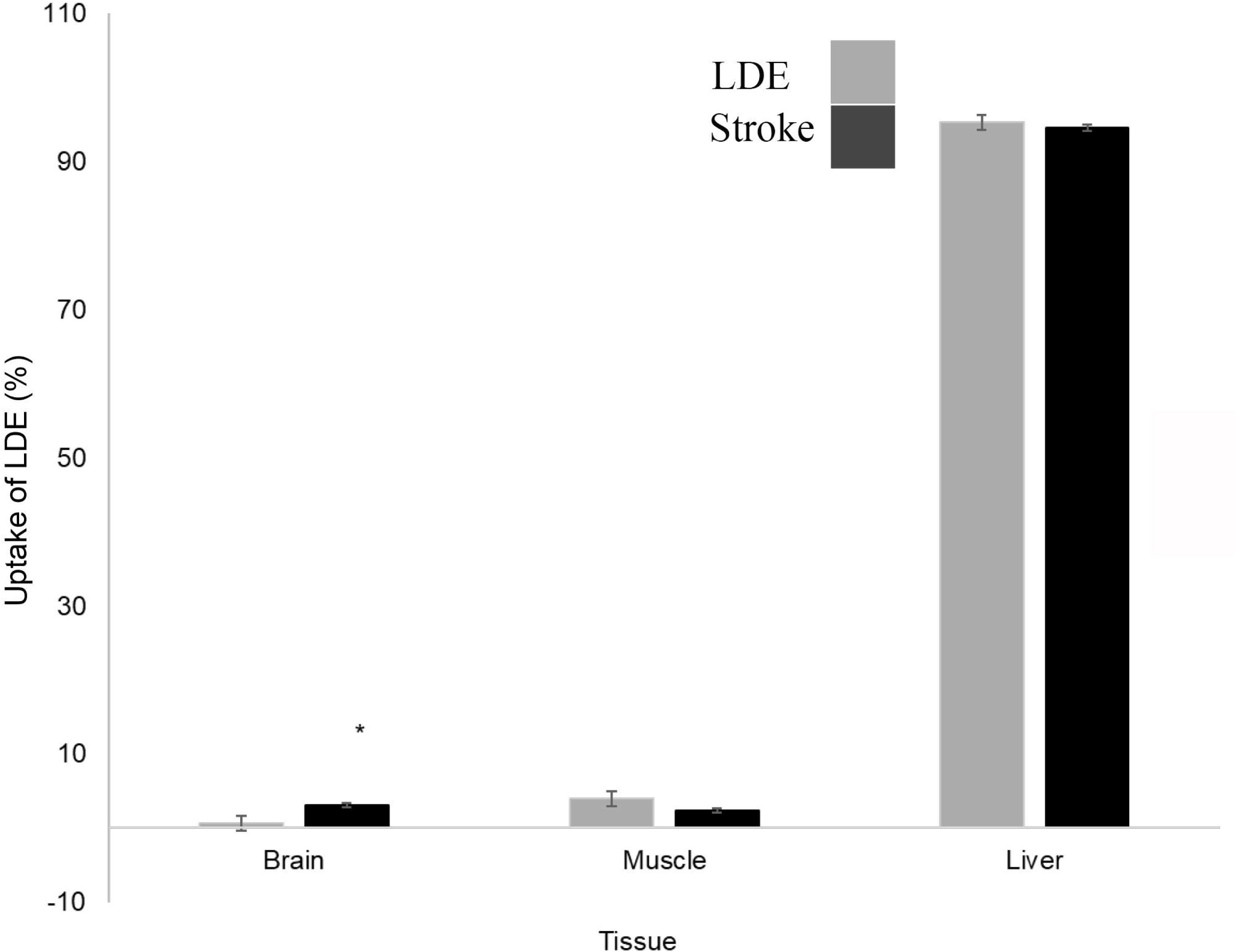
Tissue uptake assessed by scintigraphy at 24hs after intravenous injection of 3H-LDE. Comparisons between sham (n=18) and stroke (n=11) animals. Most of tissue uptake was present in the liver. Brain uptake was higher in ischemic animals compared to sham (*p<0.01). Data are expressed as normalized percentage of tissue uptake per tissue gram. Statistical comparisons were performed using unpaired Student t test with Welch’s correction.

The normalized percentage of 3H-LDE uptake per gram of tissue was higher in the brain of ischemic animals compared to control (Fig. 1 and Table-2, p<0.01). There was no uptake difference in other organs between ischemic and sham animals (Fig. 1 and Table-2, p>0.01).

### 3.2. LDE-MTX treatment reduces the inflammatory infiltrate in the ischemic core without reducing the primary infarct area

The results showed that the ischemic animals treated with unconjugated LDE presented classic focal ischemic damage, with tissue pallor and intense inflammatory response characterized by the presence of mononuclear cells in the ischemic core (Figs. 2A-B). LDE-MTX treated animals presented an almost complete abolishment of inflammatory infiltrate in the ischemic core (Figs. 2B-C). The two groups showed similar areas of primary infarct.

**Figure 2.**
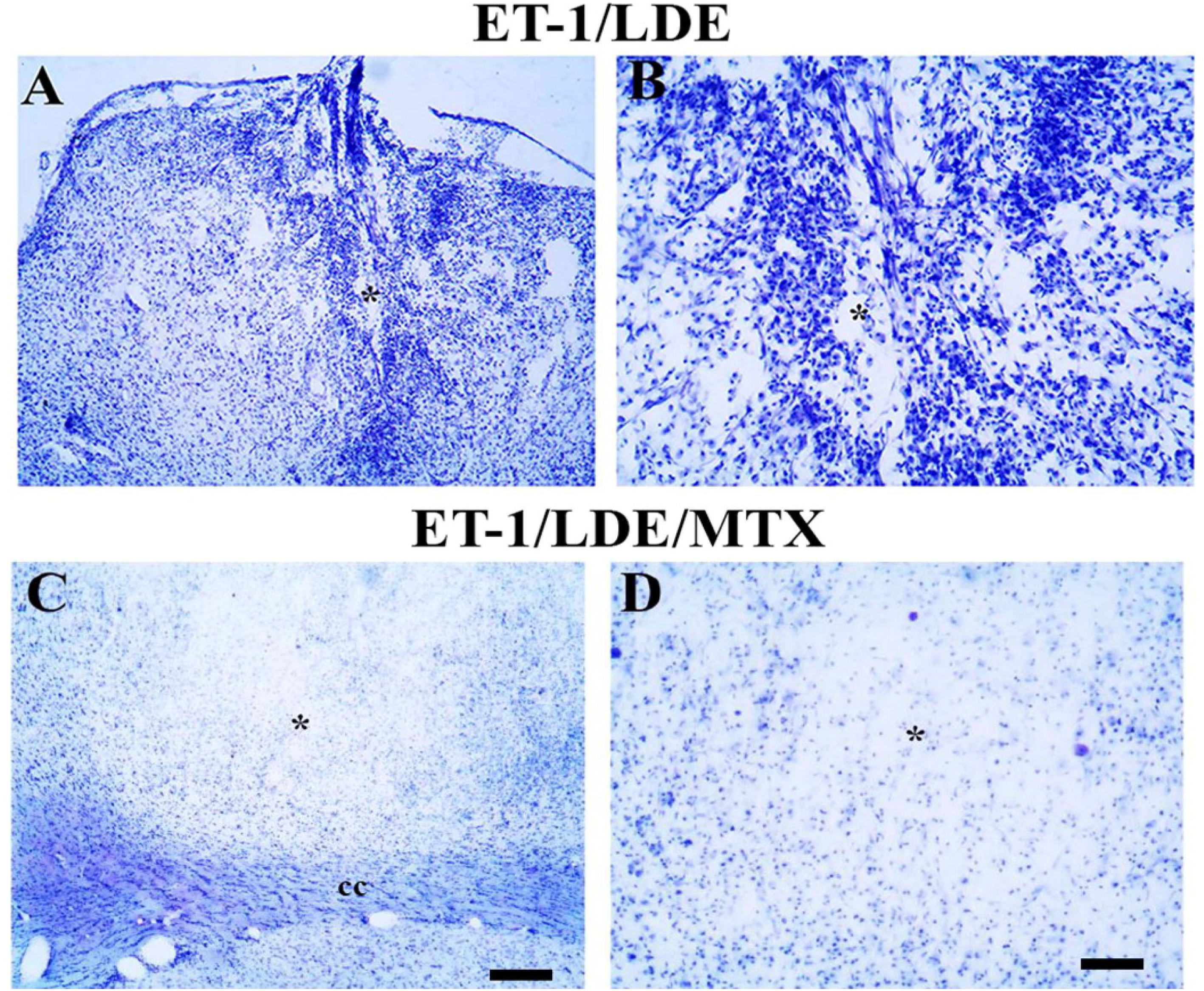
Gross histopathology using cresyl violet for ischemic animals treated with unconjugated LDE (A-B) or conjugated LDE/MTX (LDE-MTX). Infarct core was not affected in both experimental groups, but there was a conspicuous decrease in the). inflammatory infiltrate in animals treated with LDE/MTX (C-D). Asterisks point to the center of infarct core. Scale bars: A and C (200 *μ*m). B and D (80 *μ*m

### 3.3. LDE-MTX treatment reduces microglial activation after cortical ischemia

As an exacerbated microglial activation contributes to secondary damage after focal ischemia (Yrjanheikki *et al.*, 1999; Hamby *et al.*, 2007), we explored the effects of LDE-MTX treatment on microglial activation. There was intense microglial activation in animals injected with ET-1 and treated with unconjugated LDE (Figs. 3D-F). In these animals, there was a predominance of rounded microglia with phagocytic morphology at the ischemic core (Figs. 3E-F). In the periphery of the lesion, microglia with amoeboid morphology could be observed (data not shown). Ramified microglial were predominant in the contralateral side (Figs. 3A-C)

**Figure 3.**
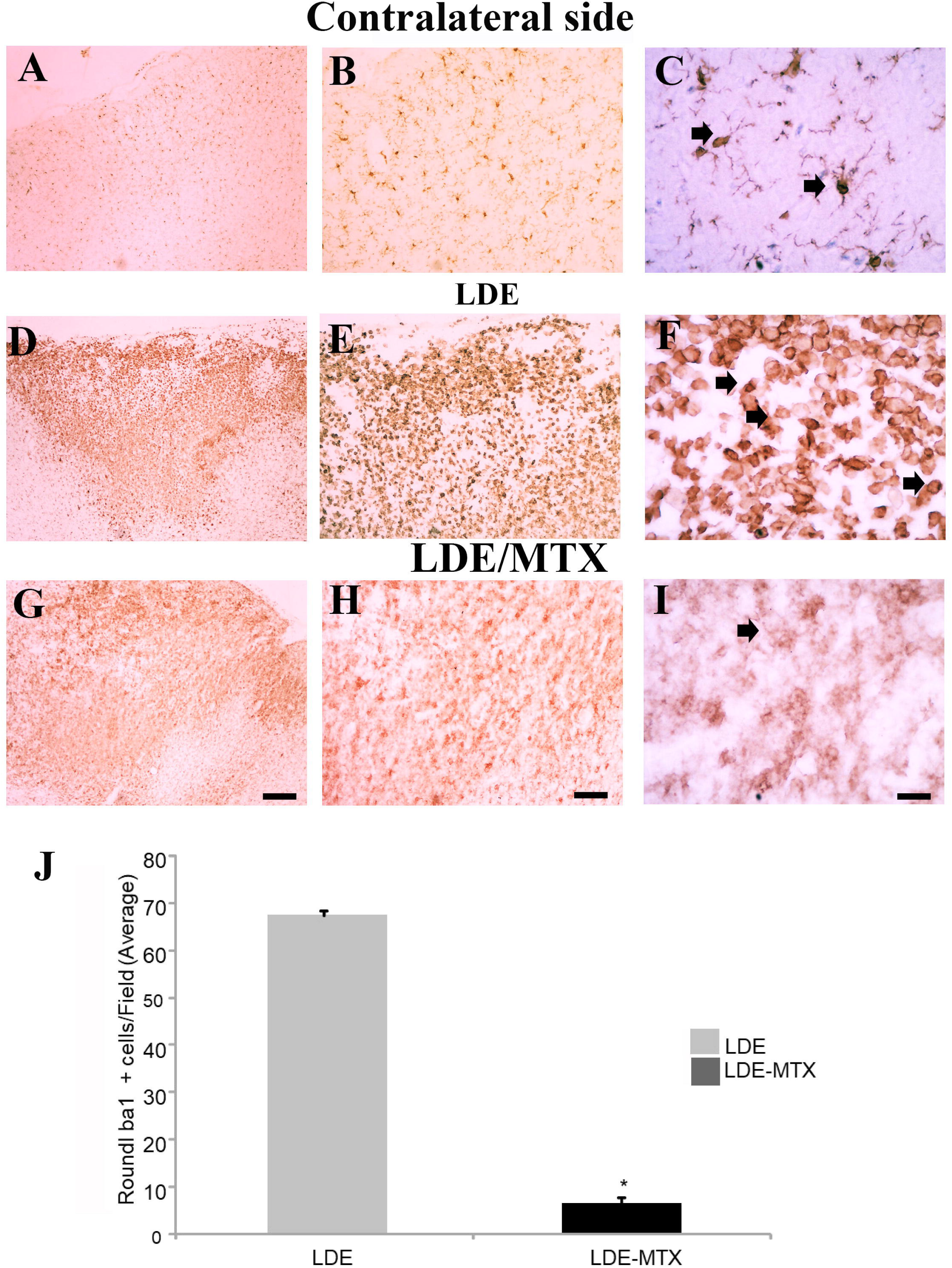
LDE-MTX treatment inhibits microglial activation (Iba1+ cells) after cortical ischemia. Resting microglia in the contralateral site (A-C) and ischemic animals treated with unconjugated LDE (D-F) or LDE-MTX (G-I) at 7 days-post injury. The LDE-MTX considerably decreases microglial activation (F). This result was confirmed by quantitative analysis of round Iba+ cells (J, * p <0.01, unpaired Student’s t test). C, F, I are higher power of B, E, H. Arrows point round Iba-1+ cells (phagocytes). Scale bars: A, D, G (200 *μ*m), B, E, H and C, F, I (80 *μ*m).

LDE-MTX treatment considerably reduced microglial activation (Fig. 3G-I). There was a great decrease in the amount of rounded microglia at the lesion center as well as amoeboid microglia around the ischemic core (Fig. 3H-J), indicating that LDE-MTX is a potent microglial inhibitor.

The qualitative results were confirmed by quantitative analysis (Fig.3J). There was a great decrease in the number of round Iba1 + cells in LDE-MTX-treated animals (6.55 ± 1.28) compared to LDE alone (67.17 ± 4.19) (Fig. 3J, p <0.01, Student t test). This corresponds to a 1025% reduction in the number of highly activated microglia.

### 3.4. LDE-MTXc treatment does not change astrocytosis after focal cortical ischemia

We addressed the effect of LDE-MTX treatment on astrocytosis. Ischemic animals treated with unconjugated LDE showed absence of astrocytes at the ischemic core (Fig. 4A) and intense astrocytosis at the periphery of the ischemic lesion (Figs. 4B-C). These activated astrocytes were present in both gray and white matters, including corpus callosum (Figs. 4B-C). A similar pattern of astrocytosis was found in animals treated with the LDE-MTX (Figs. 4D-F), indicating that MTX treatment has no effect on astrocytosis. Considering that quite similar patterns of astrocyte reactivity were found in both experimental groups, we did not quantify numbers of astrocytes.

**Figure 4.**
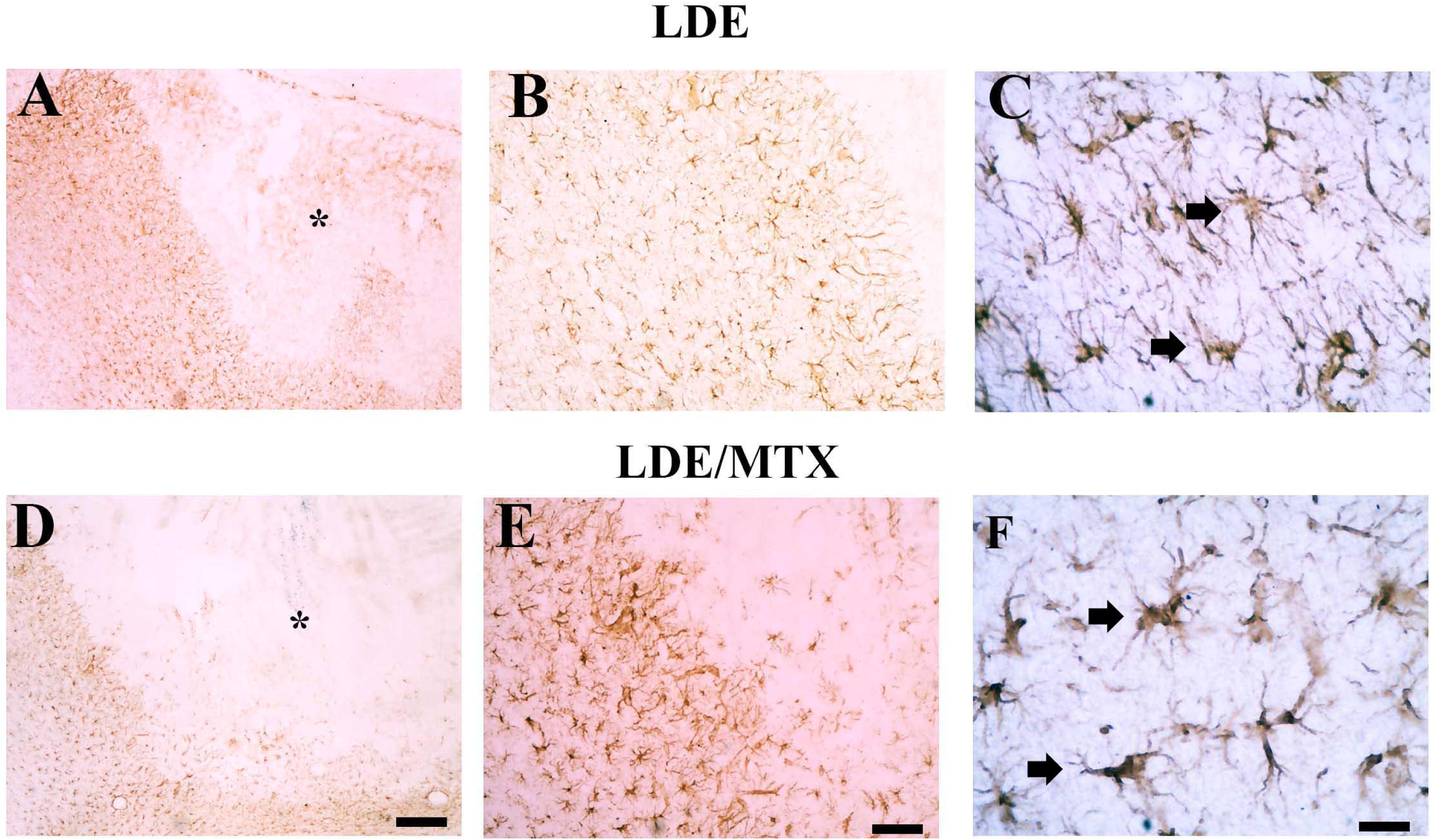
LDE-MTX treatment does not affect astrocytosis. Astrocytic activation (GFAP + cells) in ischemic animals treated with unconjugated LDE (A-C) or LDE-MTX (D-F). There was no difference on the pattern of astrocyte activation for both experimental groups. C, F are higher power of B, E. Arrows point to activated astrocytes. Scale bars: A, D (200 *μ*m), B, E (80 *μ*m) and C, F (20 *μ*m).

### 3.5. LDE-MTX induces considerable neuronal preservation in the periinfarct region

We explored the effect of LDE-MTX treatment on neuronal preservation after cortical ischemia. ET-1 microinjections into the rat motor cortex induced conspicuous neuronal loss at the ischemic core (Fig. 5A). There was no difference between areas of primary ischemic injury between LDE-MTX and LDE treated animals (Figs. 5A, D). However, there was considerable preservation of NeuN+ cell bodies in the periinfarct region in LDE-MTX treated animals (Fig. 5E-F). These results were confirmed by quantitative analysis (Fig. 5J).

**Figure 5.**
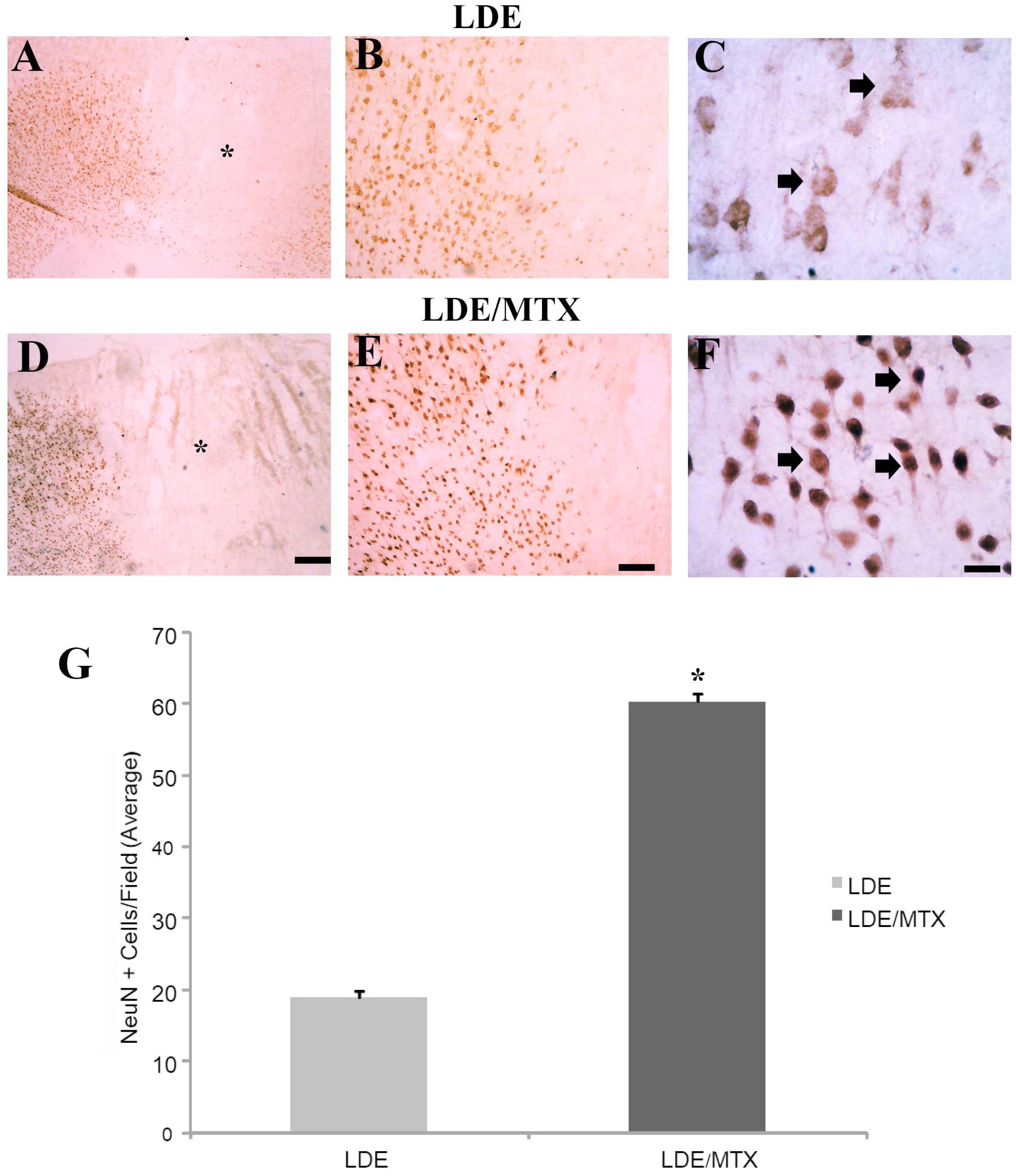
LDE-MTX is neuroprotective after cortical ischemia. NeuN + cells in ischemic animals treated with unconjugated LDE (A-C) or LDE-MTX (D-F). Intravenous treatment with LDE-MTX induce an 319% increase in neuronal preservation in the infarct periphery (D-F). This was confirmed by quantitative analysis (* p<0.01, unpaired Student’s t test). Asterisks point to ischemic lesion core. Arrows m), B, E (80 point to NeuN+ cell bodies. Scale bars: A, D (200 *μ*m), B, E (80 *μ*m) and C, F (20 *μ*m).

The quantification of NeuN+ cell bodies in the periinfarct region revealed that LDE-MTX treated group presented higher numbers of neuronal bodies per field (60.28 ± 1.37) compared to control (18.86 ± 0.88) (Fig. 5J). This data corresponds to an 319% increase in neuronal preservation, suggesting a considerable neuroprotective effect of LDE-MTX following experimental cortical ischemia.

## 4. Discussion

Stroke is a challenging health problem in several countries and an extreme cause of death and functional impairment (Iadecola *et al.*, 2011). Secondary damage is a pivotal event on the stroke pathophysiology and the main cause of functional disability (Kaiser & West, 2020; Lo, 2003; Moskowitz *et al.*, 2010). It follows that the search for effective neuroprotective agents, not yet available for human stroke, is of fundamental scientific and clinical importance.

We have investigated the anti-inflammatory and neuroprotective effects of LDE-MTX in an experimental model of cortical ischemia, as previous studies reported promising anti-inflammatory and tissue protective actions of LDE-MTX following experimental arthritis (Mello *et al.*, 2013), myocardial infarction (Maranhao *et al.*, 2017) and other diseases (Bulgarelli *et al.*, 2013; Barbieri *et al.*, 2017; Fiorelli *et al.*, 2017; Gomes *et al.*, 2018).

We first confirmed that intravenously injected LDE-MTX unequivocally crosses BBB and accumulates in different brain regions, which is a *sine qua non* condition for a good neuroprotective agent. We further demonstrated that LDE-MTX is a potent microglial inhibitor rendering a 10-fold decrease in the numbers of round activated macrophages compared to control. More importantly, LDE-MTX treatment induced a considerable 319% increase in neuronal preservation in the periinfarct region compared to control animals.

The results indicate that brain tissue bioavailability of LDE-MTX is quite expressive, considering that it was detected, without exception, in all tested animals. In addition, in pathological conditions including stroke and trauma, in which BBB rupture ocurrs, nanoemulsions like LDE may exhibit a much more significant penetration and distribution (Maranhão et al., 1993). The fact that LDE-MTX promptly crosses BBB toward the rat brain can be related to LDE diameter, chemical composition and liposolubility (Maranhão et al., 1993). The exact mechanism by which LDE crosses BBB should be addressed in future investigations.

LDE-MTX treatment does not change the primary infarct damage, but induces considerable neuroprotection in periinfarct area. Ischemic core damage is due to loss of blood flow in the central territory of vascularization of the vessels constricted by ET-1 microinjections (Fuxe *et al.*, 1989; Agnati *et al.*, 1991). Cell death and necrosis in the ischemic core occur very quickly, which renders this region normally not amenable to neuroprotection. Studies using magnetic resonance imaging (MRI) show that primary infarct area remains even in patients treated with thrombolytic therapy after ischemic stroke in humans (Lo, 2003).

Primary ischemic damage expands toward ischemic penumbra in months after ischemic stroke onset in humans (Moskowitz *et al.*, 2010). Mismatch MRI shows that this pathological expansion may increase the primate infarct area in more than 65% (Moskowitz *et al.*, 2010), which illustrates the considerable amount of salvageable neural tissue amenable to neuroprotection. The counterpart of this phenomenon in experimental models of stroke is illustrated by the progressive loss of NeuN+ cell bodies in the periinfarct area. We reported that LDE-MTX induced a 319% preservation of NeuN+ cell bodies in the periinfarct area, which suggests that MTX carried in lipidic core nanoparticles is a promising neuroprotective agent for ischemic stroke.

The exact mechanism by which LDE-MTX protects neurons in the periphery of the infarct is unknown, but this effect may be related to modulation of inflammatory response, as suggested by our previous studies in non-neural tissues (Mello *et al.*, 2013; Maranhao *et al.*, 2017). MTX is a folate analogue with antiproliferative and immunosuppressive activity used in certain types of cancer and autoimmune inflammatory diseases (Abolmaali *et al.*, 2013). The anti-inflammatory effect of MTX seems to occur through multiple actions, including folic antagonism, inhibition of eicosanoids, metalloproteinases, adenosine accumulation and blockage of lymphocyte proliferation with the corresponding production of tumor necrosis factor α (TNFα, IL-8, IL-12 and IL12, among others (Braun *et al.*, 2009).

In our previous studies, we have compared the therapeutic efficacy of LDE-MTX and commercial MTX (Mello *et al.*, 2013; Maranhao *et al.*, 2017). LDE-MTX is more effective than commercial MTX in reducing inflammatory response in an experimental model of arthritis in rabbits (Mello *et al.*, 2013). We have also shown potent anti-inflammatory and tissue protective effects of LDE-MTX in an experimental model of myocardial infarction in rats (Maranhao *et al.*, 2017). In this study, comparisons between therapeutic effects of LDE-MTX with commercial MTX revealed conspicuous differences on therapeutic efficacy.

LDE-MTX treatment was more efficacious in reducing inflammation, infarction size, myocyte hypertrophy, myocardial fibrosis, oxidative stress, cell death and improved several heart functions, including left ventricular systolic function, cardiac dilation reduction and left ventricular mass in infarcted rats (Maranhao *et al.*, 2017). In addition, it mediates increase in growth factor release in the infarcted heart contributing to both tissue protection and functional recovery.

It has been reported that commercial MTX can present serious collateral effects in some treatment schemes for certain diseases (Agarwal *et al.*, 2011; Vagace *et al.*, 2011; Salkade *et al.*, 2012; Watanabe *et al.*, 2018). A stroke-like neurotoxicity has been reported following intrathecal injections in humans (Watanabe *et al.*, 2018). This is a serious issue that could jeopardize the use of MTX as a neuroprotective agent. Nevertheless, we have previously shown that the incorporation of MTX to LDE is of fundamental importance to increase anti-inflammatory effect, tissue protection and to avoid tissue toxicity (Moura *et al.*, 2011; Barbieri *et al.*, 2017; Maranhao *et al.*, 2017; Gomes *et al.*, 2018).

The results showed that LDE-MTX considerably reduced microglial activation (a more than 10-fold decrease). This is an impressive modulation of microglial reactivity not previously afforded by any anti-inflammatory drug tested for CNS diseases. Other studies using the semi-synthetic tetracycline minocycline (Yrjanheikki *et al.*, 1999), indomethacin (Lopes *et al.*, 2016) or an inhibitor of poly (ADP-ribose) polymerase (Hamby *et al.*, 2007) have shown that inhibition of an exacerbated microglial activation induces neuroprotection. In most of these studies, similar to those reported here, reduction of primary infarct area was not observed. This has been confirmed by a recent study using the photothrombosis stroke model in sensory motor cortex of rats (Yew *et al.*, 2019).

We have reported that non-steroidal anti-inflammatory drug indomethacin decreases microglial activation after focal striatal ischemia induced by microinjections of ET-1 in adult rats (Lopes *et al.*, 2016), which was concomitant with an increase in endogenous neurogenesis and neuronal preservation without reduction of the original primary infarct area. Nevertheless, minocycline reduces infarct size in both cortex and striatum following MCAO in rats (Yrjanheikki *et al.*, 1999). We have also observed reduction of primary infarct size following ET-1 induced cortical damage into the rat motor cortex and following treatment with minocycline, bone barrow mononuclear cells or both (Franco *et al.*, 2012). Despite of this fact, reduction of microglial activation and neuronal preservation afforded by LDE-MTX treatment are much more pronounced than the observed following minocycline treatment.

It is likely that intense microglial modulation provided by LDE-MTX is fundamental for the considerable neuroprotection here reported. Although microglial cells have beneficial and neuroprotective effects after ischemia (Neumann *et al.*, 2006b; Lalancette-Hebert *et al.*, 2007; Neumann *et al.*, 2008), their detrimental effects are well described after uncontrolled activation in both acute (Yrjanheikki *et al.*, 1999; Hamby *et al.*, 2007) and chronic (Burguillos *et al.*, 2011; Sala Frigerio *et al.*, 2019) neural disorders as well as in vitro (Butovsky *et al.*, 2005; Burguillos *et al.*, 2011).

In pathological conditions, microglial cells release large amounts of proinflammatory cytokines, high mobility group box 1 (HMGB-1), nitric oxide, free radicals, proteases and other compounds that can contribute to bystander neural damage (Yrjanheikki *et al.*, 1999; Hamby *et al.*, 2007; Hayakawa *et al.*, 2008; Burguillos *et al.*, 2011). HMGB-1 is a non-histone DNA binding protein that is released in large amounts in the cellular environment after ischemic events and contributes to neuroinflammation and secondary injury (Liu *et al.*, 2007; Kim *et al.*, 2008; Muhammad *et al.*, 2008). High levels of extracellular HMGB-1 are released during ischemic injury and this inflammatory mediator is found in increased concentrations in the blood of stroke patients (Muhammad *et al.*, 2008). In addition, It has been reported that one of the mechanisms by which minocycline affords neuroprotection is by the inhibition of the extracellular release of HMGB-1 by activated microglia (Hayakawa *et al.*, 2008). It is reasonable to suggest that LDE-MTX can induce similar effects, but this should be investigated in future studies. LDE-MTX might modulate microglial activity by suppressing detrimental effects and maximizing the beneficial ones.

Although LDE-MTX conspicuously modulates microglial reactivity, there was no effect of this compound on astrocytosis. This suggests that the neuroprotective effects afforded by LDE-MTX is not related to modulation of astroglial function. This has also been shown in classical studies using minocycline (Leonardo *et al.*, 2009; Matsukawa *et al.*, 2009) and in experimental models of hemorrhagic stroke (Mestriner *et al.*, 2015), although a recent study suggests that minocycline treatment increases astrocyte activation following photothrombosis in the sensory system of adult rats contributing to neuroprotection (Yew *et al.*, 2019). These discrepant results should be related to the different experimental models used.

## 5. Conclusion

This study suggests a novel neuroprotective approach for ischemic stroke using intravenous infusion of MTX carried in lipid core nanoparticles. This compound possesses physicochemical characteristics that unequivocally enable it to promptly cross rat BBB even in the absence of brain damage, accumulating in different brain regions with no overt demise to other organs. LDE-MTX contributes to a 10-fold reduction in microglial activation affording 319% of neuronal preservation in the periinfarct ischemic area without affecting astrocytosis. As intravenous route is practical and feasible in human beings, the use of LDE-MTX must be seriously considered as a promising neuroprotective agent for ischemic stroke. This endeavor deserves future studies using different concentrations of LDE-MTX, longer survival times and primate models of stroke, before assessing the suitability for clinical trials in human.

## Acknowledgments

This study was supported by the Brazilian National Council for Scientific and Technological Development (CNPQ-Brazil). Authors are grateful to Heart Institute, University of São Paulo (Brazil), for development of LDE and LDE-MTX used in the present study.

## Data Availability Statement information

The data that support the findings of this study are available from the corresponding author, Walace Gomes-Leal, upon reasonable request.

